# Resource reallocation in engineered *Escherichia coli* strains with reduced genomes

**DOI:** 10.1101/2020.10.19.346155

**Authors:** Ernesto S. Nakayasu, Adam M. Chazin-Gray, Ryan M. Francis, Arielle M. Eaton, Deanna L. Auberry, Nathalie Muñoz, Joseph A. Cottam, Jeremy D. Zucker, Neeraj Kumar, Carrie D. Nicora, Hugh D. Mitchell, Young-Mo Kim, William C. Nelson, Robert G. Egbert

## Abstract

A major challenge in synthetic biology is properly balancing evolved and engineered functions without compromising microbial fitness. Many microbial proteins are not required for growth in regular laboratory conditions, but it is unclear what fraction of the proteome can be eliminated to increase bioproduction and maintain fitness. Here, we investigated the effects of massive genome reduction in *E. coli* on the expression level and evolutionary stability of a model biosynthetic pathway to produce the pigment protodeoxyviolacein (PDV). We identified an amino acid metabolism imbalance and compromised growth that were correlated with elimination of genes associated with significant proteome fraction. Proteomic profiling suggested that increased amino acid pools are responsible for an alleviation of fitness defects associated with PDV expression. In addition, all strains with genome reductions that significantly affected the proteome exhibited decreased stability of PDV production compared to the wild-type strain under persistent PDV expression conditions despite the alleviation of fitness defects. These findings exhibit the importance of balancing evolved functions with engineered ones to achieve an optimal balance of fitness and bioproduction.

## INTRODUCTION

Microbial bioproduction has gained interest in the last decades since it has the potential to provide a renewable source of materials, chemicals and energy (Ramos et al., 2017). Many efforts have been made to engineer organisms toward improving bioproduction (reviewed in (Chubukov et al., 2016)). It has been proposed that genomic deletion of non-essential genes could liberate cellular resources that can be directed to engineered functions (Mizoguchi et al., 2008; Reuss et al., 2017). However, a major hurdle for a more rational approach to engineering organisms is the lack of understanding of how cells reallocate cellular resources when non-essential metabolic pathways are removed.

Cells support essential processes by supplying sufficient resources for minimal maintenance, growth and reserve (Erickson et al., 2017; Metzl-Raz et al., 2017). Although maintaining a reserve of cellular resources seems to be a waste, it is essential for survival under stressful environmental conditions (Metzl-Raz et al., 2017). Among this reserve are proteins that are only required in specific environmental conditions. Therefore, many genes and their corresponding proteins are dispensable for industrial or standard lab growth conditions (Joyce et al., 2006). Genome reduction is an approach that aims to understand or improve microbial function by removing as many dispensable genes as possible without compromising growth rate.

Genome reduction has been undertaken in multiple bacterial hosts to eliminate as much as a third of the host genome. Genome reduction of the model industrial host *Bacillus subtilis* 168 yielded a strain with 7 deletions (332 genes) comprising 7.7% of its genome (Westers et al., 2003). Despite differences in cell motility, the engineered strain had minimal changes in carbon metabolism and exhibited no growth defect. Further engineering of this *B. subtilis* strain resulted in the deletion of 36% of the genome, corresponding to 2.4% of the proteome. This genome-reduced strain did exhibit a growth defect, likely as a result of reduced ribosome levels. The strain also exhibited a shift from glycolysis to an increase in amino acid metabolism^3^. In *Pseudomonas putida*, a deletion of 3.2% of the genome, including regions coding abundant proteins such as flagellar proteins, improved the production of a green fluorescent protein reporter (Lieder et al., 2015; Martinez-Garcia et al., 2014).

In *E. coli*, a library of 29 genome-reduced strains, the KHK collection, was engineered with large, sequential genomic deletions from the parental W3110 strain, including strain DGF-298 with gene deletions comprising 1.67 megabases (over 35%) of the wild-type genome (Hirokawa et al., 2013; Mizoguchi et al., 2008). While this collection of strains exhibited an increase in L-threonine bioproduction (Mizoguchi et al., 2008), a detailed growth analysis exposed a consistent reduction in growth in M63, MAA and LB media (Kurokawa et al., 2016). Choe et al. (Choe et al., 2019) found that MS56, a genome-reduced variant of *E. coli* strain MG1655, exhibited a growth defect in minimal media and imbalances in the three most important cofactors (ATP, NADH, and NADPH), as well as disrupted pyruvate and deoxynucleotide metabolism. Through multi-omics characterization of strain eMS57, which was evolved from MS56 by successive passaging in minimal media, they improved growth phenotypes and corrected much of the metabolic imbalances exhibited by MS56. These studies demonstrate that genome reduction can liberate cellular resources, but they provide limited context for the fate of those resources as they are reallocated into native and engineered metabolic pathways. They also do not provide an understanding of the benefits of genome reduction on cellular fitness and the stability of engineered functions, which represent fundamental knowledge gaps in synthetic biology research.

Here, we performed a systematic multi-omics characterization of the KHK strain collection (Hirokawa et al., 2013; Mizoguchi et al., 2008) under the burden of an engineered biosynthetic pathway to investigate the extent to which genome reduction affects the distribution and fate of metabolic resources. We used whole-genome sequencing, shotgun proteomics, and metabolomics in the presence and absence of the expression burden of an engineered protodeoxyviolacein (PDV) pathway (Cress et al., 2016). The PDV biosynthetic pathway is a truncated version (*vioABE*) of the full violacein biosynthetic pathway (*vioABCDE*) found in *Chromobacterium violaceum* (Choi et al., 2015). The PDV pathway was expressed from a pBBR1 broad-host range plasmid with tight regulation of *vioABE* by the cumate-responsive regulator CymR. The pathway utilizes the proteins VioA, VioB, and VioE, to catalyze the conversion of L-tryptophan to PDV, a dark-hued pigment with antimicrobial and anti-cancer properties (Dantas et al., 2013).

We found that large genomic deletions of dispensable genes from the KHK library liberated cellular resources related to protein translation including pools of amino acids for degradation and usage in other pathways. These large, sequential genomic deletions also led to an imbalance in glutamine metabolism, compromising cellular growth. This phenotype was alleviated by overexpressing the PDV pathway, which helped reallocate unused translational resources. The improvement in growth did not improve stability of PDV production. In fact, we found that genome reduction reduced the genetic stability of the pathway compared to the wild-type strain. These results have important implications for the development of bacterial hosts capable of performing industrial-scale, cost-competitive, and sustainable production of commodity chemicals.

## RESULTS

### Genome reduction has non-linear effects on growth and bioproduction

To start our investigation on resource reallocation, we first sequenced the genomes of the KHK strain library and compared each to the expected sequence as described in Mizoguchi et al and Hirokawa et al (Hirokawa et al., 2013; Mizoguchi et al., 2008). The genomes were assembled with a read-based genomic analysis using the published reference sequence of the parental strain, *E. coli* W3110 (Genbank accession AP009048.1). This analysis confirmed most of the expected deletions, but identified regions in deletion strains (Steps) 4, 11, 12 and 18 and MGF-01 and 02 that were still present despite being annotated as being deleted (**Figure 1A**). In Step 3, we found two deleted regions that were not expected to be deleted (**Figure 1A**). We also detected an insert in all strains, except for DGF298 (**Figure 1A**).

**Figure 1.**
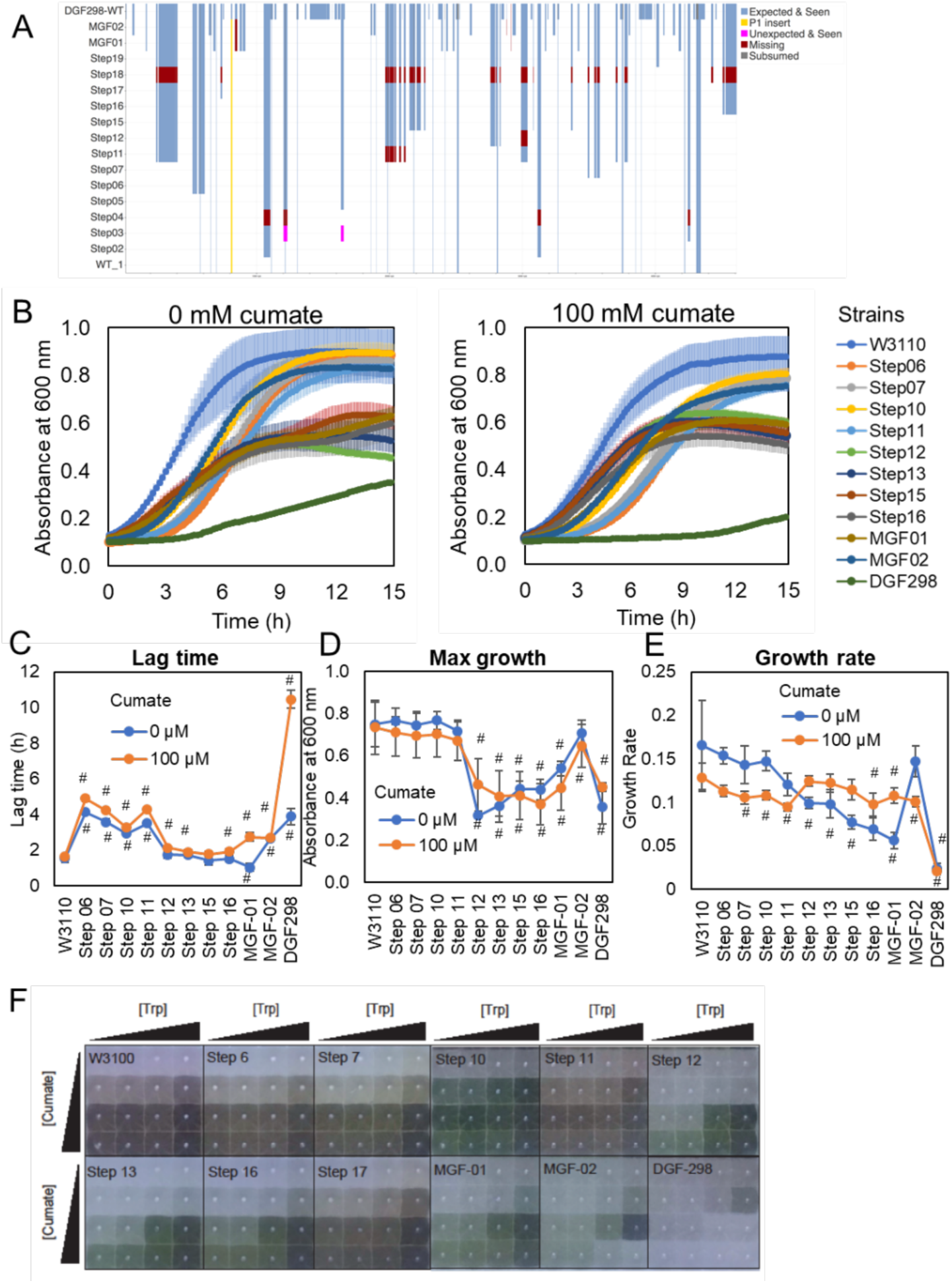
Characteristics of the engineered *E. coli* strains. (A) Genome sizes of the engineered strains. (B) Growth curves of the engineered strains with or without cumate induction of the prododeoxyviolacein pathway. (C-E) Culture lag time (C), maximum growth absorbance (D) and growth rate (E) of engineered strains. (F) Images of prododeoxyviolacein production by the engineered strains in different concentrations of the pigment precursor L-tryptophan (Trp) and expression inducer (cumate).

Next, we compared the growth of each deletion variant with and without the burden of PDV production. Growth profiles for each strain were obtained in a minimal growth medium supplemented with casamino acids with or without the addition of the inducer cumate. Strains with small genomic deletions (steps 6-11) had an approximate 2-fold increase in growth lag time compared to wild-type W3110 (**Figure 1B-C**). The longer lag time was attenuated in larger deletions (steps 12 to MGF-02) (**Figure 1B-C**). The largest deletion mutant, DGF-298 had approximately a 2-fold increase in growth lag time compared to wild-type without induction of the *vioABE* pathway and a 5-fold increase under cumate induction (**Figure 1B-C**). There was no significant effect on the growth titer of the smaller genomic deletion strains (steps 6-11) compared to W3110 (**Figure 1D**). Larger deletion strains (steps 12 to DGF-298) exhibited approximately a 2-fold decrease in the maximum growth titer in culture compared to W3110, and this was only attenuated in strain MGF-02 (**Figure 1D**). Sequential genome reduction compromised the growth rate of each strain, almost in an accumulative way, except for the outlier strain: MGF-02 (**Figure 1E**). These results suggest that genomic reduction does not reclaim cellular resources for the growth of genome-reduced strain as a linear function of genome deletion. Consistent with our initial hypothesis, the burden of *vioABE* expression reduced the growth rate of the wild-type and early-deletion strains (steps 6-11), but surprisingly increased the growth rate of the late deletion strains (step 12 to MGF-01), with the exception of MGF-02 and DGF-298 (**Figure 1E**).

To understand the impact of genome minimization on the productivity of engineered functions, we also measured the metabolic capacity of these strains to produce PDV. Steps 10 and 12 had an increase in the pigment production compared to *E. coli* W3110, which was especially noticeable at low concentrations of cumate and L-tryptophan (**Figure 1F**). Pigment production in DGF-298 was nearly imperceptible. Because of its difficulty producing pigment and its long lag phase (**Figure 1C**), we hypothesized that the induction of *vioABE* for PDV production was toxic in DGF-298. To test this hypothesis, we performed a mutant formation assay on this strain. After being grown in the presence of cumate, nearly all the DGF-298 isolates lacked pigment production (**Supplemental Figure 1**). These results showed that DGF-298 cells were extremely burdened by PDV production and that the long lag time in growth resulted in the selection of non-productive mutants.

Overall, these data showed mixed results. While some genome deletions improved bioproduction, additional deletions compromised cell growth. The reduction in growth rate is attenuated in some larger deletion strains under the burden of *vioABE* expression. These surprising results suggest that the cellular resources liberated by the large genomic deletions somehow affect the cell growth, which is ameliorated by directing these resources for PDV production.

### Genome reduction and *vioABE* expression restructure the proteome

To better understand how cellular resources are reallocated and how this affects growth and PDV production, we performed an in-depth proteomic analysis. We selected 9 of the KHK strains that represent transitions in growth or PDV production (**Figure 1**) and grew them with and without the PDV induction. The cultures were harvested in exponential growth and analyzed by liquid chromatography-tandem mass spectrometry. This analysis resulted in the identification of 2283 proteins, covering 51% of the 4451 predicted genes of the wild-type strain (W3110) containing the *vioABE* plasmid. For quantification, we used the intensity-based absolute quantification (iBAQ) method and normalized by molecular mass to calculate the fraction that each protein represented compared to the total proteome mass (**Table S1**). Our analysis confirmed that many of the deleted regions had trace or undetectable levels of protein expression (e.g. steps 1, 13, and 17) (**Figure 2A**). Thus, much of the genomic reduction of the KHK library had little effect on liberating translational resources. However, two deletion steps accounted for the largest reductions in proteome mass fraction. The regions deleted in step 12 and DGF-298 resulted in the largest proteome reductions, representing respectively 1.0 and 2.0% of the protein mass of the wild-type strain (**Figure 2A**). **Figures 2B and 2C** show a drastic reduction in protein abundance of genes deleted in step 11 and MGF-02, respectively.

**Figure 2.**
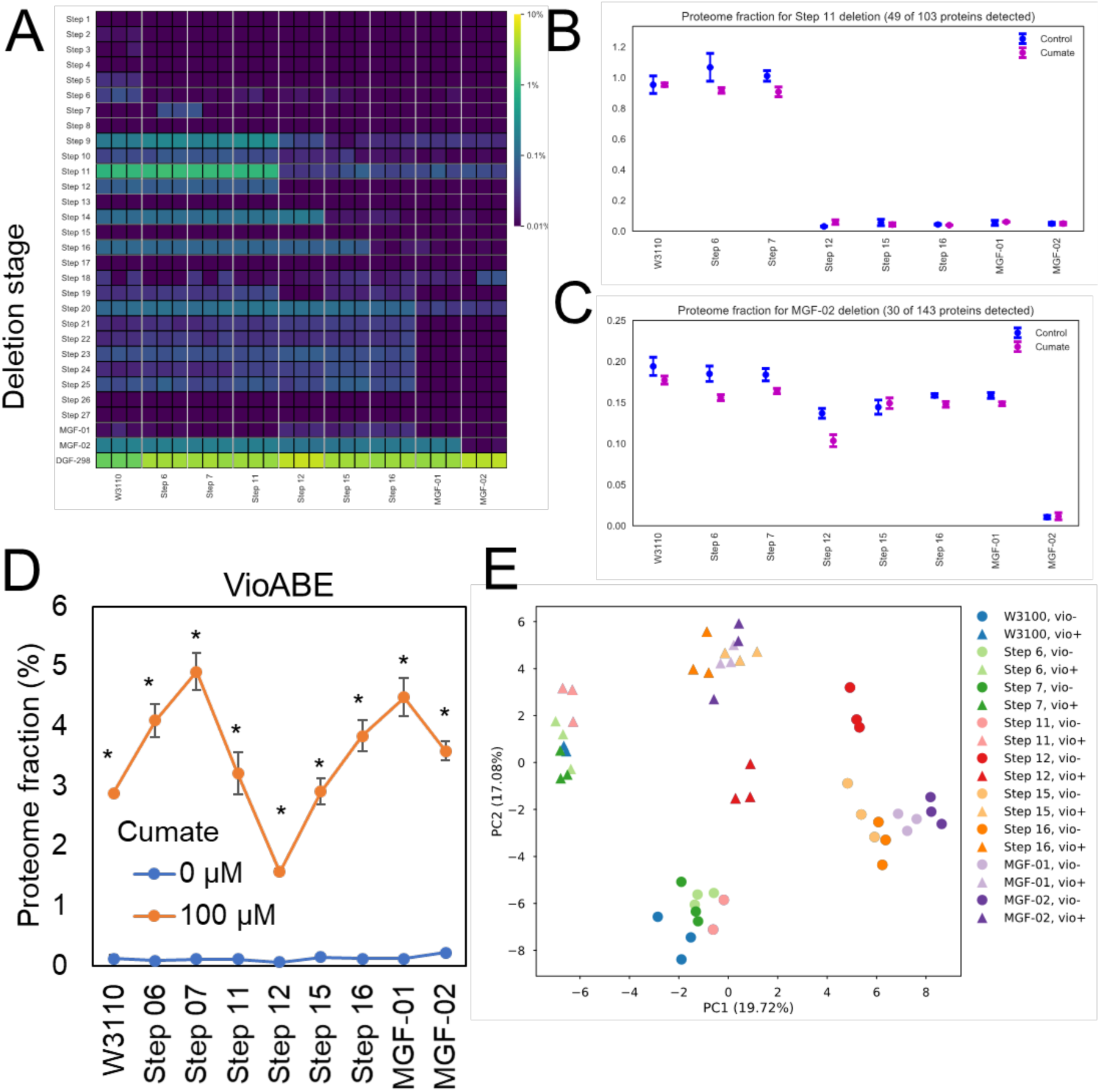
Characteristics of the proteome of the engineered strains. (A) Fraction of the proteome deleted in each of the engineered strains. (B-C) Cumulative abundance of proteins from genes deleted in step 11 (B) and MGF-02 (C). (D) Cumulative abundance profile of proteins from the protodeoxyviolacein (vioABE) pathway in the presence or absence of cumate (expression inducer). (E) Principal component analysis of the proteomics data from the different engineered strains. * p≤0.05 by T-test comparing the same strain with or without the cumate treatment.

The impact of genome reduction and PDV pathway induction on the proteome was also investigated in our analysis. As expected, the addition of cumate significantly induced expression of VioA, VioB and VioE, which collectively represented up to approximately 5% of the proteome mass in some strains (**Figure 2D**). Also, genome reduction had a non-linear positive affect on VioABE expression with pathway induction (**Figure 2D**). Both genome deletions and pigment overproduction induced significant changes to the proteome, as observed can be seen by the separation of the proteomic profiles of these strains on the principal component analysis (PCA) plot (**Figure 2E**).

### Genome deletions have no effect on cell growth related proteins

Growth rate analysis indicated that while genomic reduction negatively impacts growth rate for strains with significantly reduced genomes, it is partially restored by overexpression of *vioABE*. Also, ribosome levels have been shown to be regulated by resource availability and directly proportional to the growth rate of cells in multiple organisms (Klumpp et al., 2013; Metzl-Raz et al., 2017). Therefore, we queried the proteomics data to investigate levels of proteins belonging to pathways related to cell growth to determine if this effect could be observed in our data.

For the KHK deletion strains, the ribosome levels remained stable and was unaffected by the induction of the PDV pathway (**Figure 3A**), resulting in poor correlation between growth rate and ribosome abundance (**Figure 3B**). This relationship is linear until step 11, showing proportional growth reduction according to genome deletion size and PDV production (**Figure 3C**). We observed only minor changes in the levels of other pathways related to cell growth: cell division, RNA polymerase and DNA polymerase III complexes (**Figure 3D-F**).

**Figure 3.**
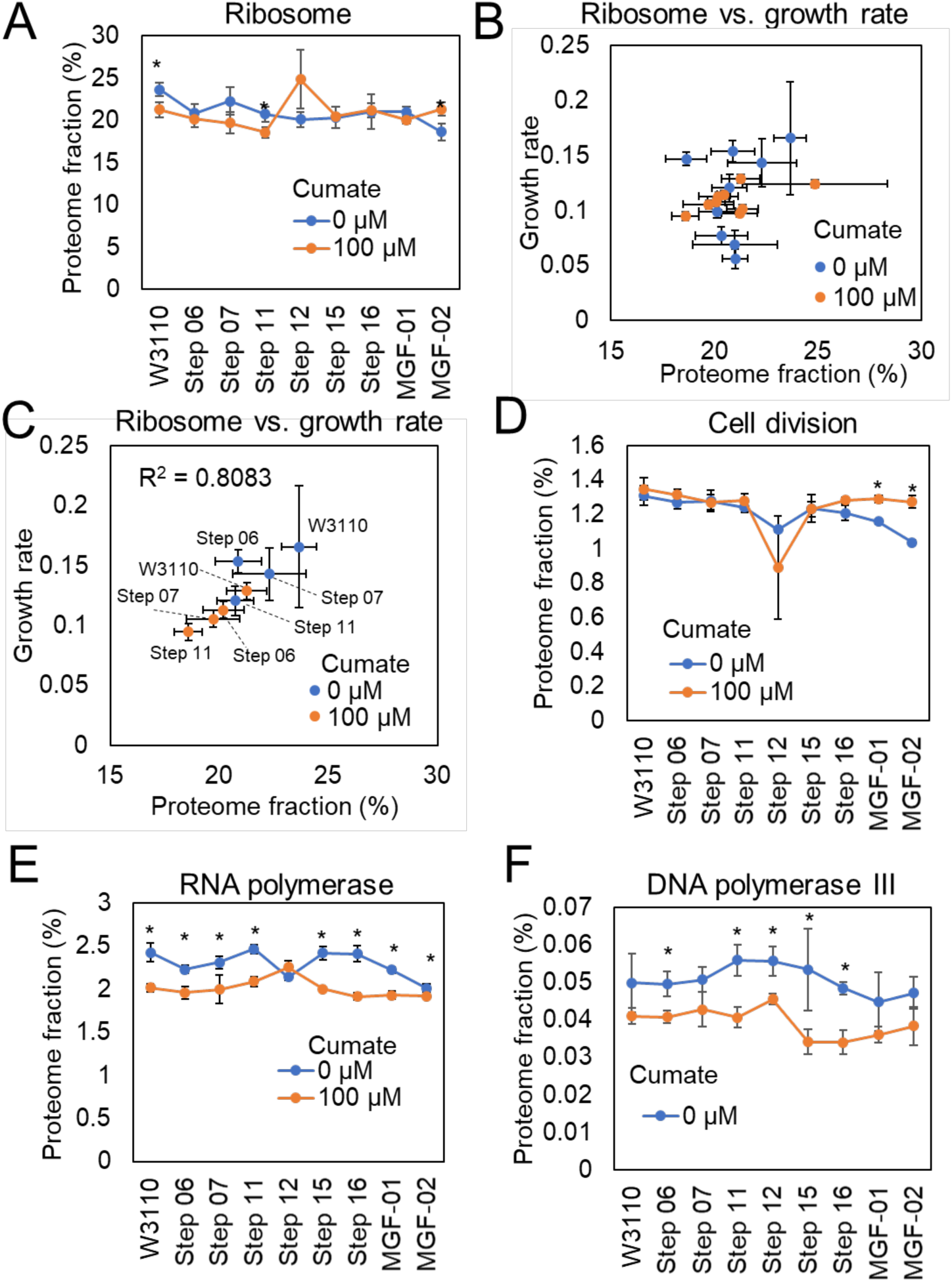
Abundance profiles of pathways directly involved in cell growth. (A) Cumulative abundance (sum of the abundance of all proteins in the pathway) profile of ribosomes. (B) Relationship between the ribosomal content of the cell and growth rate of strains in panel (A). (C) Relationship between the ribosomal content of the cell and growth rate from wild-type W3110 stain to step 11. (D-F) Cumulative abundance profile of cell division proteins (D), DNA-directed RNA polymerase complex (E) and DNA polymerase III complex (F). For expression profiles of individual proteins and comparison with the validation dataset, see figure S1. * p≤0.05 by T-test comparing the same strain with or without the cumate treatment.

To validate these findings we performed another round of proteomic analysis for strains W3110 (WT), step 12, MGF-02, and DGF298 (**Table S2**). A strikingly similar and stable expression pattern was observed for the proteins related to ribosomes, cell division and RNA polymerase and DNA polymerase III complexes in the second data set (**Figure S1**). None of the changes in these cellular pathways were correlated with growth rate alterations, suggesting that the growth defects of the engineered strains are not due to these processes.

### Amino acid metabolic pathways are downregulated by genomic reduction and *vioABE* induction

Aiming to better understand the phenotypes carried by our engineered strains, we analyzed the proteomics data to investigate how genomic reduction affects global metabolic pathways. We performed clustering based on the abundance profile followed by a function-enrichment analysis. This analysis showed a strong enrichment for proteins related to amino acid metabolism among the proteins that increase in abundance during genome reduction (**Figure 4A**). A manual inspection of impacted pathways showed a common dysfunction in glutamine metabolism in genome-reduced strains, which is summarized in **Figure 4B**. We observed a strong increase in expression of enzymes that degrade glutamine into succinate and glyoxylate. This suggests that a reduced demand for amino acids in genome-reduced strains leads to reallocation of these resource to other metabolic processes. This upregulation of glutamine catabolism was accompanied by a change in the polyamine biosynthesis pathway towards the production of putrescine, spermidine and spermine (**Figure 4B, Figure S2**).

**Figure 4.**
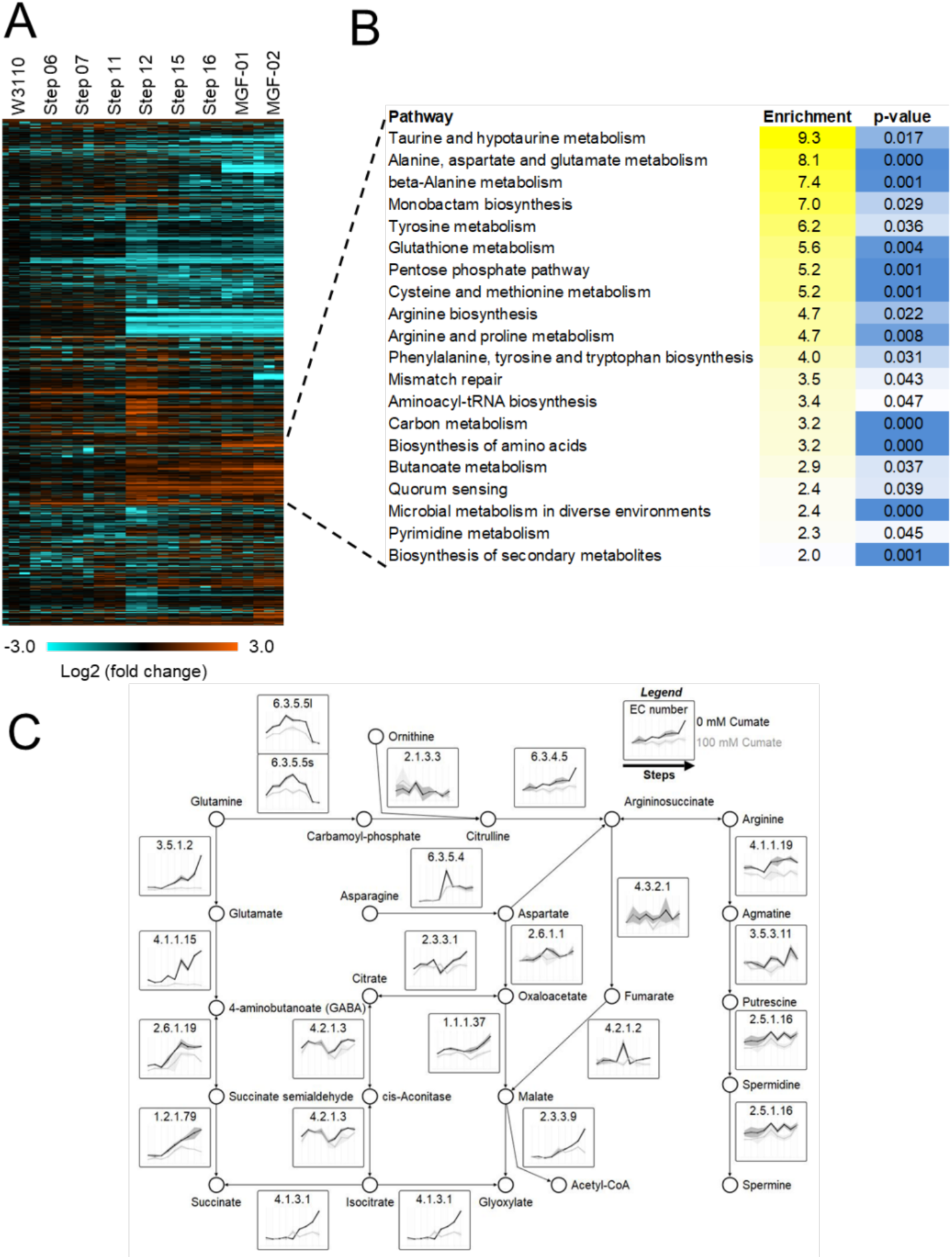
Global expression proteomic profile and pathway analysis of the engineered *E. coli* strains. (A) Heatmap of the differentially abundant protein between the engineered strains – ANOVA test, p ≤ 0.05. (B) Function-enrichment analysis of proteins of which abundances increased according to the genome deletion size. (C) Curated pathway of proteins abundances increased according to the genome deletion size. The pathway shows the degradation of glutamine into polyamines and succinate. For the full names of individual proteins, expression profiles and comparison with the validation dataset, see Figure S2.

Due to the strong upregulation of proteins in the glutamine degradation pathway, we had a closer look at other proteins that regulate glutamine levels (**Figure 5A**). The levels of glutaminase GlsA1 increase 12.5-fold from wild-type to MGF-02 (**Figure 5A-B, S3**). The levels of glutamine transporter subunits GlnQ and GlnH have a slight increase from wild-type to step 11, followed by an approximate tenfold decrease in their abundance (**Figure 5A and 5C-D, S3**). Both trends in degradation (increase) and transport machinery(decrease) are reversed by inducing burden on the cells via overexpression of the vioABE system (**Figure 5B-D, S3**), which requires a large amount of resources for pigment production. These results indicate that genome-reduced strains are trying to reduce a possible excess in glutamine levels by increasing degradation and blocking transportation unless burdened by the induction of PDV biosynthesis.

**Figure 5.**
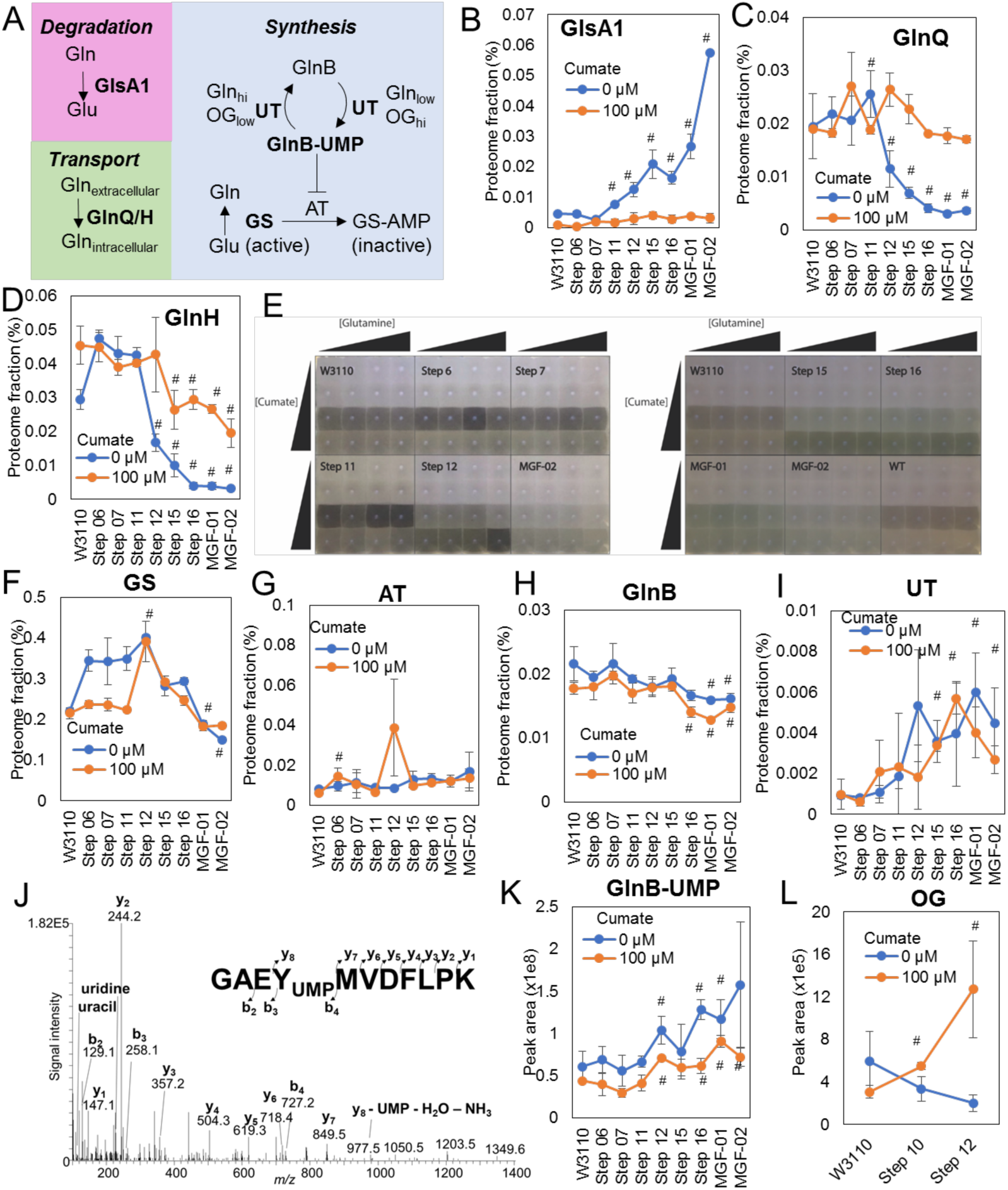
Glutamine metabolism in engineered *E. coli* strains. (A) Glutamine metabolic pathway. (B-D) Abundance profiles of glutaminase GlsA1 (B) and glutamine transporter subunits GlnQ (C) and GlnH (D). (E) Prododeoxyviolacein pigment production in strains supplemented with glutamine. (F-G) Abundance profiles of glutamine synthase GS (F), GS adenylyltransferase AT (G), nitrogen regulatory protein GlnB (H), GlnB uridylyltransferase UT (I). (J) Tandem mass spectrum of GlnB uridylylation at tyrosine 51. (K-L) Abundance profiles of uridylylated GlnB (GlnB-UMP) (K) and oxoglutarate (OG) (L). For expression profiles of individual proteins and comparison with the validation dataset, see figure S3. # p≤0.05 by T-test comparing the engineered strain against the wild type W3110 strain.

We next investigated the glutamine synthesis pathway. Nitrogen regulatory protein P-II 1 (GlnB) is UMPylated by uridylyltransferase (UT) in the presence of low glutamine concentration and high oxoglutarate (OG). UMPylated GlnB (GlnB-UMP) in turn blocks adenylyltransferase (AT), which inhibits glutamine synthetase (GS) activity by adding an adenosine monophosphate (AMP) moiety (**Figure 5A**). Therefore, under low glutamine and high OG concentrations, this pathway induces GS activity. In contrast, when there is a high glutamine and low OG concentration, GS activity is inhibited. GS levels exhibited a small, 1.8-fold increase in strains up to step 12 and a gradual reduction to basal levels by strain MGF-02 (**Figure 5E**). AT and GlnB levels remained mostly stable in all the deletion strains (**Figure 5F-G, S3**), which is consistent with previous results showing that the expression of these proteins is not affected by nitrogen levels (van Heeswijk et al., 1993).

The levels of UT increased gradually to over 6 folds in MGF-01 compared to W3110 strain (**Figure 5H, S3**), despite the fact that its expression is not regulated by nitrogen levels (van Heeswijk et al., 1993). We quantified the levels of GlnB-UMP by extracting the peak area of the modified peptide G_48_AEY_UMP_MVDFLPK_58_ (**Figure 5I, S3**) and normalizing it by the unmodified peptide I_91_FVFDVAR_98_. The level of GlnB-UMP increases by 2.6 in MGF-02 compared to the W3110 strain (**Figure 5J**), showing a positive correlation between UT and GlnB-UMP levels (R^2^ = 0.6037). Because OG regulates UT activity, we measured the levels of this metabolite by gas chromatography-mass spectrometry. However, the levels of OG had no significant changes in steps 10 and 12 compared to the W3110 (**Figure 5K**). Overall, these results showed an increase in GlnB-UMP levels in *E. coli* strains with large genomic deletions. GlnB-UMP increase is correlated with the activation of GS. Therefore, both glutamine synthesis and degradation are activated at the same time in *E. coli* strains with large genomic deletions. This indicates that genome reduction causes a general dysfunction in glutamine metabolism, which we hypothesize may play a role in growth defect observed in large deletion strains.

### Genome reduced strains exhibited decreased stability of PDV expression

To understand the balance between proteome recovery and evolutionary stability, we collected pigment expression data for a subset of the KHK library in five passages representing over ∼45 generations. Each strain was passaged independently with or without cumate induction in minimal media. At each step, an outgrowth was performed in rich media with cumate to visualize and score pigmentation **(Fig 6A)**. Although no genome-reduced strains matched the robust pathway stability seen in W3110, Step 12 notably maintained PDV production in approximately 80% of samples. Step 10 and MGF-01 performed poorly with less than 40% of cultures maintaining the pathway under prolonged induction **(Fig 6B)**. Uninduced cultures (left side of plate, **Fig 6A**) maintained pigment production in outgrowth assays for nearly all cases.

**Figure 6.**
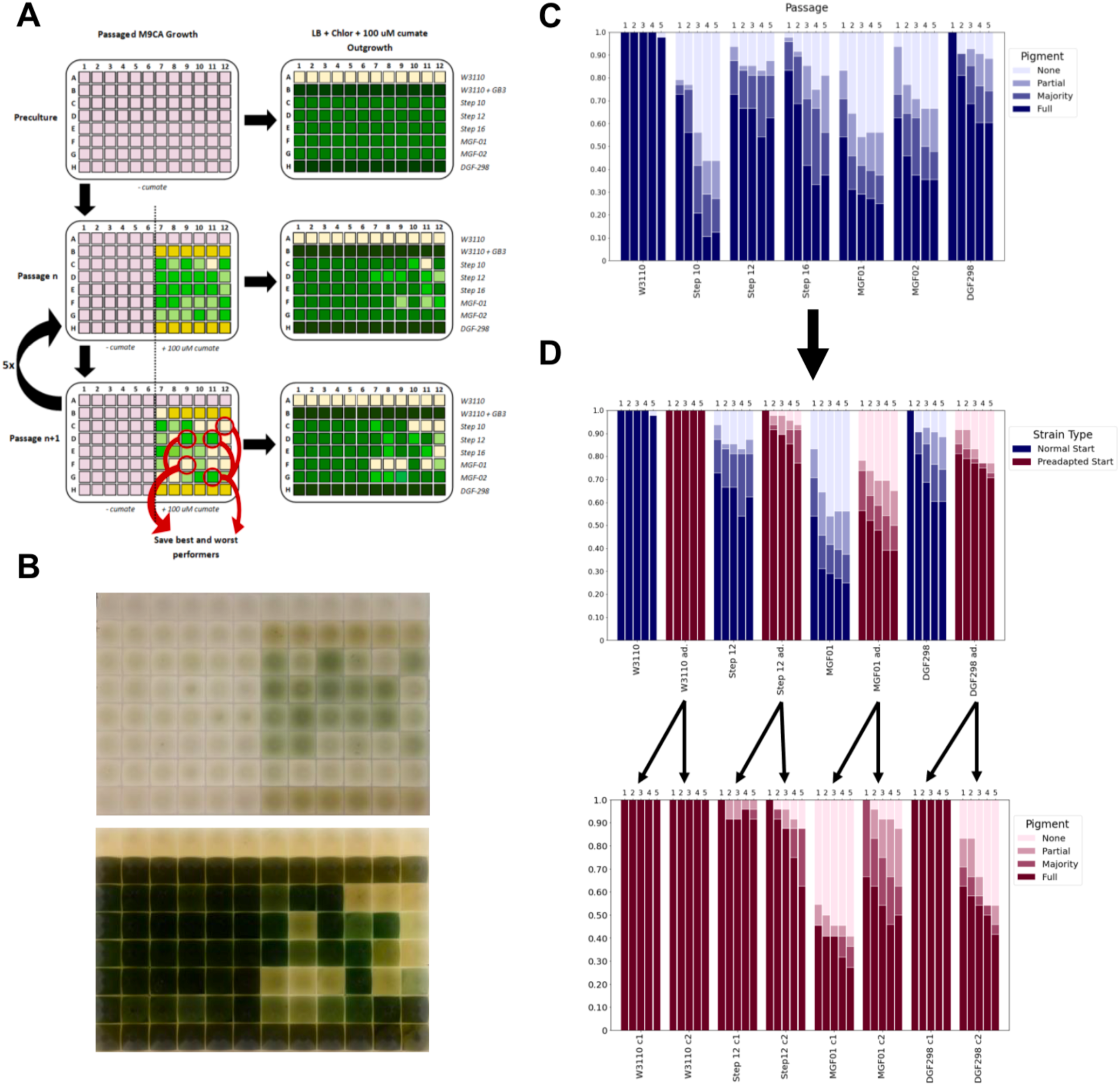
Culture passaging to visualize pathway stability over many generations. **A**. After a preculture in supplemented M9CA, cells were passaged into fresh media containing 0 or 100 µM cumate for 5 consecutive days. Cells were additionally inoculated into LB + Chlor34 + 100 µM cumate to visualize pathway retention. After the final passage, top and bottom performers, determined by pigmentation, were reserved. **B**. Example plate images. (Left) Typical M9CA passage with right half of plate under cumate induction. (Right) LB outgrowth under cumate induction. **C**. Violacein expression by strain observed from LB outgrowth after each 24-hour passage under induction in M9CA + Chlor34 + 100 µM cumate. Bars represent N = 42 cell lines divided among 5 growth plates with height reported as a fraction of N. The darkest color indicates strongly pigmented cells while the lightest indicates cells similar to a violacein-lacking W3110 control. **D**. Blue bars represent violacein expression in preadapted cells (5 passages under cumate induction) observed from LB outgrowth after each 24-hour passage under induction in M9CA + Chlor34 + 100 µM cumate. Analogous bars from subfigure C are reproduced in red. Blue bars in the upper plot represent N = 48 cell lines derived from 2 preadapted candidates for each strain with height reported as a fraction of N. Each strain-candidate is resolved on the lower plot, N = 24. The darkest color indicates strongly pigmented cells while the lightest indicates cells similar to a violacein-lacking W3110 control.

Upon completion of 5 passages, a subset of pigmented and non-pigmented (when available) cultures for each strain were saved as ‘adapted’ cultures. An identical assay was performed on these isolates to determine whether the end point of the prior induction could serve as a more stable starting point for additional experiments. While the gains were modest, we observed increased stability in both Step 12 and MGF-01. Interestingly, when data was resolved to individual isolates from the original assay (represented by red circles in **Fig 6A**, it became clear that while some isolates had no improvement compared to the starter population, others showed distinct adaptations for improved pigment stability.

Growth curves for the PDV expression stability assay revealed that growth adaptations under induced conditions were independent of pigment production. In the first passage, all strains showed some increase in lag time and/or decrease in growth rate under cumate induction. The growth profiles were consistent with earlier experiments (**Fig. 1**). After the first passage, however, a large recovery was observed, with most strains matching or nearly matching the growth profile of W3110. Additionally, this growth recovery appeared to be completely independent of engineered pathway function—i.e. pigmented and unpigmented strains grew at identical rates.

## DISCUSSION

In this paper, we investigated the reallocation of cellular resources in *E. coli* strains with large genomic deletions. We first analyzed the growth properties of these strains in M9 medium supplemented with casamino acids under no induction burden, which exposed significant growth defects for many of these strains. This growth defect was unlikely to be due to regulation of pathways related to cell proliferation, such as ribosomes, cell division proteins, DNA-directed RNA polymerase complex and DNA polymerase III complex. Although these findings are inconsistent with the results from Mizoguchi et al., subsequent work supports our conclusion that large genomic deletions do compromise growth phenotypes (Hirokawa et al., 2013; Kurokawa et al., 2016; Mizoguchi et al., 2008). This inconsistency may be a result of a difference in media composition or instability caused by the maintenance of the synthetic load plasmid. Furthermore, the observed growth defect could be the result of some toxic byproduct or an imbalance in the metabolism of strains with large genomic deletions.

In support of the hypothesis that a toxic metabolite may be affecting growth of genome-reduced strains, we found that the growth defect of strains with large genomic deletions was diminished when burdened with the overexpression of the PDV. The drain of resources by induction of the *vioABE* pathway might have alleviated the excessive buildup of metabolites like glutamine or other imbalances in metabolism. Mizoguchi et al. found a 2.4-fold increase in the production of L-threonine by the MGF-01 strain (Mizoguchi et al., 2008), which is inconsistent with our observations for PDV production by this strain. This contradictory phenotype might also be related to differences in media composition, plasmid maintenance, the buildup of toxic byproducts, or an imbalance in metabolism. Our analysis showed that genomic reduction enhanced the amino acid degradation and polyamine synthesis pathways, likely due to resources liberated by the massive genomic deletions. A deeper investigation of these pathways led to the discovery of an imbalance in the metabolism of glutamine in these strains. We found that glutaminase expression was upregulated and the glutamine transporter subunits were highly downregulated, as genomic reduction resulted in a build-up of unused glutamine.

Despite the increased availability of glutamine as a result of genomic deletions, the glutamine synthase regulatory network was significantly upregulated in larger deletion strains. Both factors likely influence the disruption in glutamine metabolism, resulting in a growth defect and consequently a reduction in PDV pigment production. We hypothesize that this metabolic imbalance can be reduced by adaptive laboratory evolution (ALE), which could aid in the acquisition of additional mutations to rebalance cell metabolism (Choe et al., 2019). In fact, this is exactly what we found after successively passaging our genome reduced strains in minimal media prior to PDV pathway induction. Growth rates of strains with large genomic deletions improved significantly after just one passage. This in turn led to increased evolutionary stability of our biosynthetic pathway in pre-adapted genome-reduced strains.

Genetic reduction diminishes the cellular demand for amino acids, reallocating them to other metabolic processes. However, extremely large genomic deletions impair the ability of the cell to regulate amino acid metabolism causing a defect in growth and bioproduction, suggesting that deletions need to be balanced by adding functions or ALE (Choe et al., 2019). Furthermore, large genomic deletions significantly reduced the evolutionary stability of our biosynthetic pathway possibly due to the deletion of genes that are non-essential for growth but that are essential for genome integrity. Therefore, we suggest that a more elaborate engineering strategy is needed to enhance the production of biocommodities without causing a disruption in metabolism, growth, or evolutionary stability. One such genome remodeling approach is called ReProMin. ReMinPro relies on a combination of gene essentiality data, transcription factor regulatory data, and proteomic abundance data to identify dispensable transcription factors. Specifically, ReProMin utilizes transcription factor regulatory data to link highly expressed nonessential genes to their corresponding regulatory genes. By knocking out transcription factors that drive expression of highly-expressed, nonessential genes, they were able to free up over 0.5% of the proteome budget without inducing a significant growth defect (Lastiri-Pancardo et al., 2020).

These findings open new perspectives for cellular capacity optimization to enhance the production of biocommodities. We anticipate genome remodeling efforts guided by multi-omics data will elucidate how the reclamation of protein expression capacity can enhance the expected behaviors and increase evolutionary stability of engineered functions, and provide novel biocontainment strategies for microbes deployed in industrial, medical, and environmental settings.

## STAR METHODS

### CONTACT FOR REAGENT AND RESOURCE SHARING

Further information and requests for resources and reagents should be directed to and will be fulfilled by the Lead Contact, Robert Egbert.

## EXPERIMENTAL MODEL AND SUBJECT DETAILS

### Bacteria strains

The plasmid cloning was performed in *E. coli* MG1655. The P1 transductions, electrocompetent cell preparations, plasmid transformations, growth, and proteomics analysis were performed in *E. coli* W3110 (and KHK derivative strains).

## METHOD DETAILS

### Growth conditions and media

*E. coli* strains were grown in LB Miller broth (10 g/L tryptone, 10 g/L NaCl, 5 g/L yeast extract) at 37 °C with shaking at 250 rpm, and supplemented with 50 µg/mL kanamycin or 34 µg/mL chloramphenicol where applicable, unless stated otherwise. For proteomics analysis, cultures of each KHK strain were inoculated from glycerol stocks into LB media supplemented with 34 µg/mL chloramphenicol. These cultures were back diluted 100-fold into M9 medium supplemented with 0.01% casamino acids, 34 µg/mL chloramphenicol, 5 µg/mL tryptophan, and 10 µg/mL biotin and grown for 12 h at 37 °C with shaking at 700 rpm. Preinduced cultures were diluted 1000-fold into 5 mL fresh media with cumate (100 µM) and without cumate and grown under the same conditions to an optical density (600 nm) of between 0.2 and 0.4. Cells were harvested by centrifuging at 7,000 x g for 10 min at 4 °C, decanting media supernatant, and storing pelleted cells at −80 °C until processing for downstream analysis.

### Sequencing and comparative analysis

KHK library (including MGF-02 and DGF-298) strains were grown to saturation in LB medium and genomic DNA was prepared from these cultures using the DNA Blood and Tissue kit (Qiagen). Extracted DNA was sent for commercial amplicon library preparation and whole genome sequencing (SNPsaurus, Eugene, OR). The breseq analysis program (Deatherage and Barrick, 2014) was used to align the resulting reads against the *E. coli* W3110 genome (Genbank accession AP009048.1) (Hayashi et al., 2006). Breseq identifies single nucleotide polymorphisms (SNPs), short deletions, and novel junctions which might imply either genome rearrangements or long deletions. The lists of deleted regions for each sequenced strain were compared against the published lists of deletions and scored as either verified, intact, or unintended.

### Plasmid cloning (pBHV-CymR-vioABE-cmR)

A chloramphenicol-resistant broad host range plasmid was constructed to introduce an inducible system of high-burden PDV production into the KHK strains (**Figure S4**). Standard Q5 Hot Start High-Fidelity Master Mix (New England Biolabs M0494S) was used to amplify the pBBR1 backbone from the pBHV-KanR plasmid (P1) using oligonucleotide primers O1 (RP GG_J23150) and O2 (pBHV_vioA_GG_fwd). The same backbone was used to amplify the vioABE genes, while introducing the promoter-repressor system pCym-CymR, using primers O3 (FP GG_CymR) and O4 (vioABE-pBHV_GG_rev). Successful amplification of these fragments was confirmed by 1% agarose gel electrophoresis. These fragments were then purified using the DNA Clean and Concentrator Kit-5 (Zymo Research). The purified amplicons were then fused by performing a Golden Gate assembly reaction (Weber et al., 2011), which was set up using 12.5 nM of the backbone amplicon, 25 nM of the vioABE-CymR insert and 25 nM of a synthesized product: O11 (BCD8). After performing a golden gate assembly reaction in a thermocycler, pBHV-CymR-KanR-vioABE-1 (P2) was transformed into Turbo electrocompetent *E. coli* (New England Biolabs) forming strain S1, which was grown to saturation, and then miniprepped (Qiagen). To retune the RBS’s of the *vioABE* genes to induce a greater burden on cells during induction of the pathway, a combinatorial library of *E. coli* strains with different strength RBS’s for a genomically inserted *vioABE* pathway was surveyed to identify a high-burden variant (Lee et al., 2013). To swap out the high-burden vioABE pathway, the pBHV backbone was amplified from plasmid P2 using primers O5 (vioSwap_pBHV_fwd) and O6 (vioSwap_pBHV_rev). Next, the vioABE genes with retuned RBS’s were amplified from the selected high-burden strain S2, using primers O7 (vioSwap_vioABE_fwd) and O8 (vioSwap_vioABE_rev). After these amplicons were run on a 1% agarose gel to confirm expected band lengths, they were PCR purified and assembled using a NEBuilder HiFi DNA Assembly (New England Biolabs) protocol, forming plasmid pBHV-CymR-KanR-vioABE-2 (P3). Next, the chloramphenicol-resistance marker was replaced for the kanamycin-resistance marker. The pBHV-CymR-vioABE-2 was transformed into the *E. coli* strain RE1000 forming strain S3. Next this strain was made electrocompetent for recombineering as described below. After performing a standard PCR amplification of the CmR gene from *E. coli* strain S4 (bRE004) using primers O9 (cmR_pBHV_fwd), and O10 (cmR_pBHV_rev), this amplicon was PCR purified, and then 300 ng were transformed into strain S3. After selecting on LB supplemented with chloramphenicol, and a culture of strain S5 (bPNL037) was grown and then mini-prepped (Qiagen) isolating the pBHV-CymR-vioABE-cmR (P4) plasmid. The sequence of this plasmid was confirmed using Nextera sample prep, followed by Illumina MiSeq sequencing and alignment analysis.

### P1 Transduction

*E. coli* KHK strains were inoculated from glycerol stocks into 3 mL of LB medium and grown overnight at 37 °C with shaking at 250 rpm. 1.5 mL of each saturated culture were transferred to a 1.5-mL microcentrifuge tube and centrifuged at 7000 x g for 2 minutes at 23 °C. After discarding the supernatant from each tube, the pelleted cells were resuspended in 750 µL of P1 salts buffer (10 mM MgCl_2_, 5 mM CaCl_2_ in water). In a separate 1.5-mL microcentrifuge tube, 100 µL of the recipient strain was resuspended in P1 salts solution was combined with 100 µL of donor phage lysate with a P1-compatible KanR cassette. This mixture was incubated at 37 °C for 30 minutes without shaking. Next, 1 mL of LB medium and 200 µL 1 M sodium citrate were added to this tube to arrest phage adsorption. This mixture was then incubated at 37 °C for 1 hour without shaking. Then, the tubes were centrifuged at 23 °C for 2 minutes at 7,000 x g. Cell pellets were resuspended in 100 µL of LB and seeded on LB agar plates supplemented with 50 µg/mL kanamycin and 5 mM sodium citrate. These plates were incubated at 37 °C for 12-16 hours before colonies were picked into LB medium and glycerol stocks were prepared.

### DGF298 sacB-cmR removal

To transform the strain DGF-298 with the pBHV-CymR-vioABE-cmR plasmid, a sacB-cmR casette that was used to make the final edits to this strain, had to be removed. Whole genome-sequencing of DGF-298 revealed the insertion location of the sacB-cmR cassette on its genome in between genes ydgL and gudP. Primers O12 (ydgL_BsaI_fwd)_and O13 (mltA_rev)_were designed to amplify the region just upstream of the cassette and O14 (gudP_fwd)_and O15 (gudP_BsaI_rev) were designed to amplify the region just downstream of the cassette. Next, after verification of expected band lengths on a 1% agarose gel, and PCR purification of these amplicons, a BsaI-golden gate reaction (New England Biolabs) was setup as described by NEB to fuse these *ydgL* and *gudP* amplicons. Successful ligation of the gudP and ydgL amplicons was confirmed on a 1% agarose gel. After a 0.4x SPRI Bead cleanup (Ampure) of the BsaI golden gate product, a nested amplicon was amplified by using a set of nested primers O16 (gupP-nested_fwd) and O17 (ydgL-nested_rev). After undergoing a second 0.4x Spri bead cleanup, the PCR amplicon was quantified on a Nanodrop (Thermo Fisher). DGF-298 electrocompetent cells were prepared for recombineering as previously described except that at an optical density (600 nm) of 0.2 the 50 mL culture of DGF-298 was spiked with 100 ng/mL of anhydrotetracycline (aTc) and grown to a final optical density (600 nm) of 0.4-0.6. DGF-298 cells were washed twice with 10 mL of a chilled 10% glycerol solution and were resuspended in 100 µL of 10% glycerol. DGF-298 electrocompetent cells were transformed with 300 ng of the purified PCR product (described above) by electroporation in a Genepulser XCell (Biorad) at 1800 V, 25 µF, and 200 Ω in a 1-mm gap length electrocuvette (Bulldog Bio). After recovering for 2-3 hours in 300 µL of pre-warmed SOC at 37 °C, 150 µL of culture was seeded on LB plates supplemented with 5% sucrose to select against the retention of the SacB-cmR cassette. After confirming the removal of the sacB-cmR cassette by PCR, gel electrophoresis, and testing sensitivity to chloramphenicol, colonies were picked into LB medium, grown to saturation at 37 °C with 250 rpm of shaking, and glycerol stocks were stored at −80 °C.

### Transformations

Electrocompetent cells were prepared as previously described^22^. Aliquots of each strain were thawed on ice for 10 minutes before being mixed with 100 ng of pBHV-cymR-vioABE-CmR purified plasmid. This mixture was then transferred into a 2-mm gap length 96-well electroporation plate (HT-P96-2, Harvard Apparatus) and electroporated in a Gene Pulser XCell (Biorad) at 2500 V, 25 µF, and 200 Ω. Electroporated cells were immediately transferred into pre-warmed SOC recovery media (New England Biolabs) and recovered for 1-2 hours at 37 °C with 250 rpm of shaking. After recovery, each culture was plated onto LB agar supplemented with chloramphenicol and grown at 37 °C for 24 hours. Colonies were picked into and grown in liquid LB media supplemented with 34 µg/mL chloramphenicol for 24 hours at 37 °C with 250 rpm of shaking. Glycerol stocks of each strain were made by mixing an equivalent volume of 50% glycerol and saturated culture, and stored at −80 °C.

### Echo inoculation and microplate reader growth analysis

KHK strains were inoculated from glycerol stocks into LB medium containing 34 µg/mL chloramphenicol and grown overnight at 37 °C with 250 rpm of shaking. Saturated starter cultures were diluted 100-fold into M9 medium supplemented with 0.01% casamino acids, 34 µg/mL chloramphenicol, 5 µg/mL tryptophan and 10 µg/mL biotin, and grown for 12 h at 37 °C with shaking at 700 rpm. M9 pre-induced cultures were diluted 5-fold into sterile nuclease-free water and 50 µL of each was loaded into an Echo Acoustic Liquid Handler (Labcyte) compatible 384-well source plate (Labcyte PP-0200). A 40 mM cumate solution (40% ethanol) was loaded into the same source plate. After preparing the source plate, a Greiner 384-well destination plate (Sigma-Aldrich M1811) was filled with 80 µL of the supplemented M9 medium. An echo transfer sheet was generated using a Python program (See Supplemental Code I) that created an array of 12 strains, 4 cumate conditions, and 8 replicates of each condition. This transfer sheet was uploaded into and run through the Echo Plate Reformat software (Labcyte). After liquid transfer was completed by the Echo, the destination plate was covered with a Breathe-easy seal (Diversified Biotech) and transferred to a Synergy H1 Microplate Reader (Biotek). A kinetic run was then initiated for 24 hours at 37 °C, shaking continuously at maximum speed in a double orbital configuration. The plate reader was programmed to collect absorbance (600 nm), and fluorescence (Ex: 585 nm, Em: 625 nm) measurements of every well in 10-minute intervals. Custom Python code was then used to analyze and plot this kinetic growth and fluorescence data.

### Proteomics analysis

Pelleted cells were extracted, and proteins were digested as previously described (Nakayasu et al., 2017). Digested samples were dissolved in water and randomized prior to data acquisition on the LC-MS/MS system (Yang et al., 2017). Briefly, 500 ng of peptides were loaded into a trap column (5 cm x 360 µm OD x 150 µm ID fused silica capillary tubing (Polymicro, Phoenix, AZ); packed with 3.6-µm Aeries C18 particles (Phenomenex, Torrance, CA). Peptides were separated in a capillary column (70 cm x 360 µm OD x 75 µm ID packed with 3-µm Jupiter C18 stationary phase (Phenomenex) with a gradient of acetonitrile in water containing 0.1% formic acid. The eluate was analyzed online in a quadrupole-orbitrap mass spectrometer (Q-Exactive Plus, Thermo Fisher Scientific, San Jose, CA). High-energy collision dissociation was performed for the 12 most intense ions, before dynamic excluding them for 30 sec.

### Genetic stability of PDV pathway

A subset of the complete KHK library (W3110-WT, W3110+GB3, Step 10, Step 12, Step 16, MGF-01, MGF-02, DGF298) was streaked onto LB supplemented with 34 μg/mL chloramphenicol and 10 µM cumate, and grown overnight at 37°C. 10 µM cumate was used to screen for contaminants and to pick only pigment-producing colonies at the first step. This low-level induction was sufficient to select candidates without introducing undue burden to the cells. Dark colonies were picked from each plate into 1 mL of fresh M9 medium supplemented with 0.01% casamino acids, 34 µg/mL chloramphenicol, 5 µg/mL tryptophan and 10 µg/mL biotin and grown overnight at 37°C in a tabletop shaker at 1000 rpm. Saturated cultures were inoculated at a ratio of 1:500 into 1 mL of fresh media with the addition of either 100 or 0 µM cumate and grown under identical conditions. At inoculation, 200 μL of culture from each well was added to a Corning 3795 96-well flat-bottom plate and OD_600_ data were collected every 10 minutes until saturation in a Biotek H1 plate reader.

Cells were passaged a total of 5 times in this way. At each step, cultures were also inoculated into 1 mL of fresh LB supplemented wtih 34 µg/mL chloramphenicol and 100 µM cumate, and grown under the same conditions as above. Images were taken through the bottom of each LB plate by shining a diffuse white light through the top of the plate into a lightbox. A Nikon D7500 camera was used to collect images.

## QUANTIFICATION AND STATISTICAL ANALYSIS

### Proteomics quantification

Collected LC-MS/MS proteomics data were analyzed with MaxQuant software (v.1.5.5.1 and v.1.6.5.0) (Tyanova et al., 2016). Peptide identification was performed by searching against the in-lab sequenced version of the W3110 genome containing the sequences for all used plasmids. Only fully tryptic peptides were considered for the searches and methionine oxidation was set as a variable modification. Parent and fragment mass tolerances were set as the default parameters of the software. Identifications were filtered at 1% false-discovery rate at both peptide-spectrum match and protein levels. Proteins were quantified using the intensity-based absolute quantification (iBAQ) method (Sandberg et al., 2019). For absolute quantification, iBAQ values of individual proteins were normalized by the protein molecular mass and then by the total mass in the sample, resulting in relative protein mass of the cell. GlnB-UMP was identified and quantified manually using QualBrowser (Xcalibur v4.0.27.42). Peaks were integrated and smoothed with Gausian 13 before extracting their areas. Proteins were considered differentially abundant using ANOVA test.

## Acknowledgments

This work was funded by PNNL’s Agile Investment in Synthetic Biology. Samples were analyzed in the Environmental Molecular Sciences Laboratory, a DOE BER national scientific user facility on the PNNL campus. PNNL is a multi-program national laboratory operated by Battelle for the DOE under contract DE-AC05-76RL01830.

## AUTHOR CONTRIBUTIONS

ESN, ACG, RE conceived the study and designed the experiments; ACG, RMF, AME, NMM, CN and RE performed the experiments; ESN, ACG, RMF, AME, DLA, NMM, JAC, JDZ, NK, HDM, YMK, WCN and RE analyzed the data; and ESN, ACG, RMF, NMM, and RE wrote the manuscript with inputs from all the authors. All authors read and approved the final manuscript.

## Data availability

The proteomics data were deposited in Pride repository (Perez-Riverol et al., 2019), a member of the ProteomeXchange consortium (Vizcaino et al., 2014).

Set 1 – Project accession: PXD016365

Username

Password: kVYtATEr

Set 2 - Project accession: PXD016380 Reviewer account details:

Username

Password: 2bcW7JuQ

---

**Figure S1.**
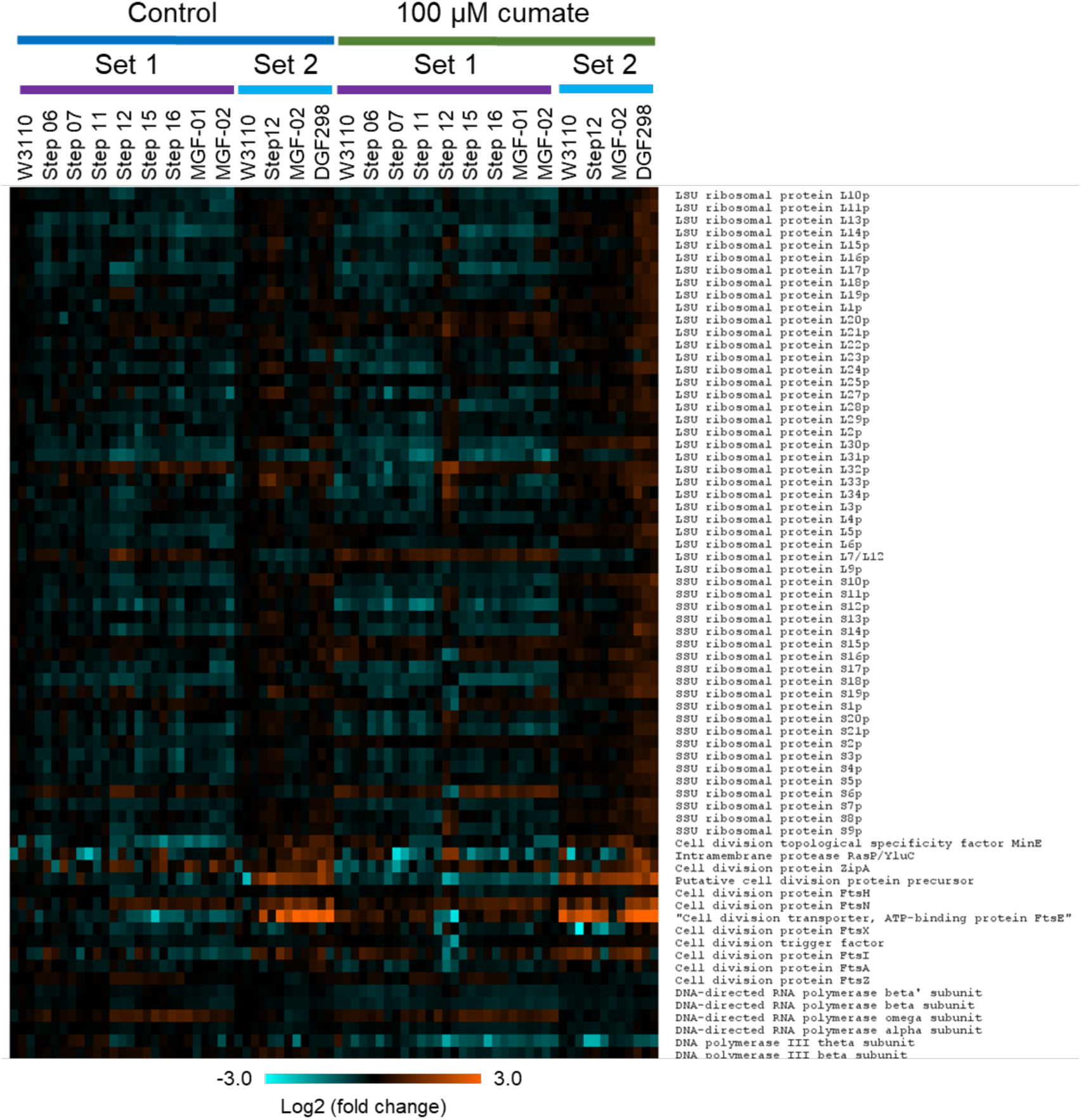
Expression profiles of individual proteins from pathways directly involved in cell growth (Figure 3). Proteins from the ribosome, cell division, DNA-directed RNA polymerase and DNA polymerase III pathways are shown in the heatmap. Proteomics analysis was carried out in two sets, for discovery (set 1) and validation (set 2) purposes. The engineered strains were grown in M9 medium with casamino acids with or without 100 μM cumate to induce expression of the *vioABE* pathway.

**Figure S2.**
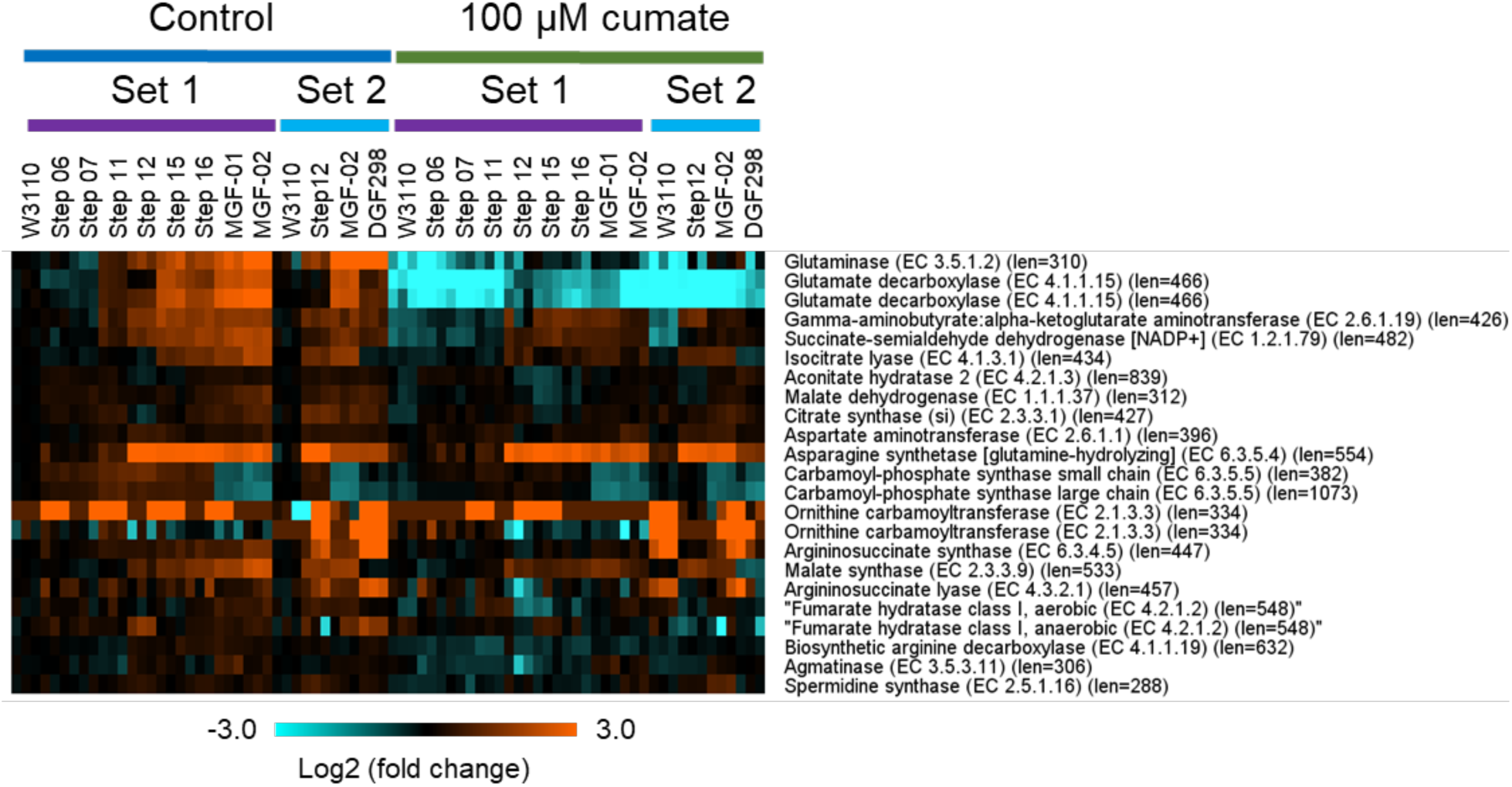
Expression profiles of individual proteins involved in the degradation of glutamine (Figure 4). Proteomics analysis was carried out in two sets, for discovery (set 1) and validation (set 2) purposes. The engineered strains were grown in M9 medium with casamino acids with or without 100 μM cumate to induce expression of the *vioABE* pathway.

**Figure S3.**
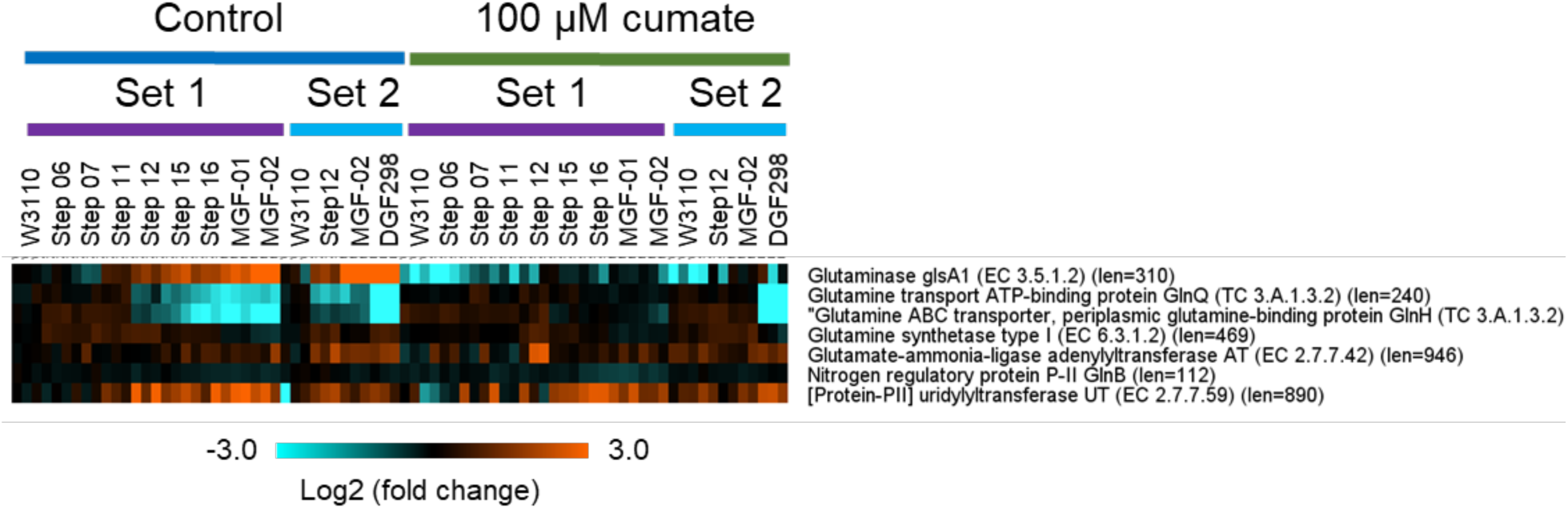
Expression profiles of individual proteins involved in the glutamine regulation (Figure 5). Proteomics analysis was carried out in two sets, for discovery (set 1) and validation (set 2) purposes. The engineered strains were grown in M9 medium with casamino acids with or without 100 μM cumate to induce expression of the *vioABE* pathway.

**Figure S4.**
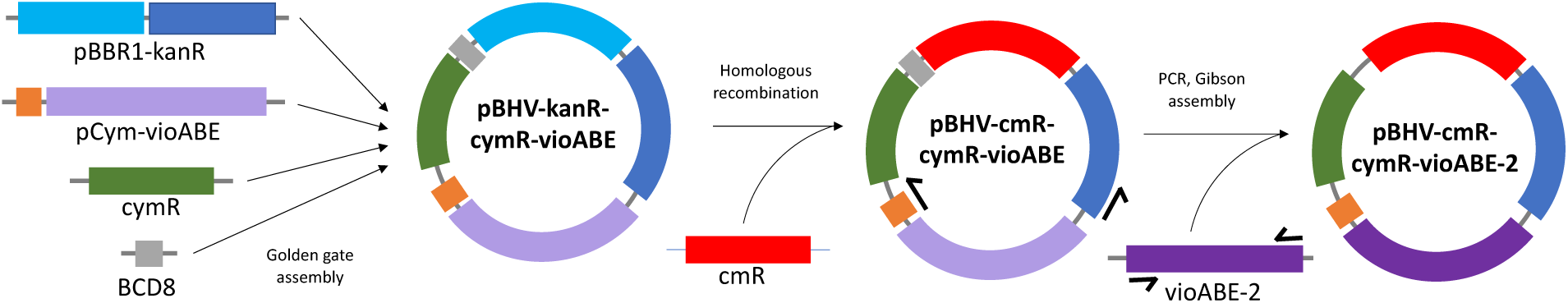
Overview of plasmid cloning schema. A broad-host range, kanamycin-resistant (kanR) plasmid backbone was initially assembled with a pCym/CymR regulated vioABE pathway. Next, the kanamycin-resistance marker was replaced with a chloramphenicol-resistance marker (cmR). Finally, the vioABE genes were PCR amplified off of a high burden variant from a vioABE RBS library and assembled with the chloramphenicol-resistant plasmid backbone to form the final plasmid – pBHV-cmR-cymR-vioABE-2.

